# Monitoring the 5’UTR landscape reveals 5’terminal oligopyrimidine (TOP) motif switches to drive translational efficiencies

**DOI:** 10.1101/2021.07.02.450886

**Authors:** Ramona Weber, Umesh Ghoshdastider, Daniel Spies, Clara Duré, Fabiola Valdivia-Francia, Merima Forny, Mark Ormiston, Peter F. Renz, David Taborsky, Merve Yigit, Homare Yamahachi, Ataman Sendoel

## Abstract

Transcriptional and translational control are key determinants of gene expression, however, to what extent these two processes can be collectively coordinated is still poorly understood. Here we use long-read sequencing to document the 5’and 3’untranslated region (UTR) isoform landscape of epidermal stem cells, wild-type keratinocytes and squamous cell carcinomas. Focusing on squamous cell carcinomas, we show that a small cohort of genes with alternative 5’UTR isoforms exhibit overall increased translational efficiencies and are enriched in ribosomal proteins and splicing factors. These 5’UTR isoforms with identical coding sequences either include or exclude 5’terminal oligopyrimidine (TOP) motifs and result in vastly altered translational efficiencies of the mRNA. Our findings suggest that switching between TOP and non-TOP motif-containing 5’UTR isoforms is an elegant and simple way to alter protein synthesis rates, set their sensitivity to the mTORC1-dependent nutrient-sensing pathway and direct the translational potential of an mRNA by the precise 5’UTR sequence.

## Introduction

Gene expression is tightly regulated in space and time to determine cellular function and behavior. Transcriptional and translational control are key steps in the gene expression pathway; however, to what extent these two processes can be collectively coordinated is still poorly understood. Recently, several studies have suggested a non-canonical mode of regulation, by which a large cohort of yeast genes switch between short and long 5’ untranslated regions (UTRs) while keeping the coding sequences (CDSes) identical (Chen et al., 2017; Cheng et al., 2018; Hollerer et al., 2019; Tresenrider et al., 2021). Since 5’UTRs are critical for ribosome recruitment and ultimately initiation of translation (Hinnebusch et al., 2016), switching between these 5’UTR isoforms resulted in differential translational efficiencies of these mRNAs. During yeast meiosis, for instance, around 8% of genes were found to be temporally regulated by an extended 5’UTR that was poorly translated due to the presence of inhibitory upstream open reading frames (uORFs) (Cheng et al., 2018). These examples indicate the possibility that the presence or absence of regulatory sequences set by the exact 5’UTR isoform could directly control protein synthesis rates of an mRNA.

In this study, we used nanopore long-read sequencing to document the UTR isoform landscape of *in vivo* epidermal stem cells, wild-type keratinocytes and cultured squamous cell carcinoma cells (SCC^c^). We identified 496 and 243 significantly altered alternative splicing events in squamous cell carcinomas and epidermal stem cells, respectively. Focusing on squamous cell carcinomas, we show that the subset of genes with altered 5’UTR isoform usage in SCC^c^ exhibit overall increased translational efficiencies and are enriched in ribosomal proteins and splicing factors. Interestingly, we observed that for two examples of ribosomal protein transcripts, *Rpl21* and *Rpl29*, SCC^c^ switch to a different set of 5’UTR isoforms that either expose or mask 5’terminal oligopyrimidine (TOP) motifs. TOP motif-containing mRNAs are targeted by the mammalian target of rapamycin complex 1 (mTORC1) nutrient-sensing pathway to selectively enhance their translation rates (Hsieh et al., 2012; Thoreen et al., 2012).

Our findings suggest that switching between TOP and non-TOP motif-containing 5’UTR isoforms is a previously unappreciated, elegantly simple way to effectively alter translational efficiencies of a cohort of genes in squamous cell carcinomas. More generally, since alternative transcription start sites (Pelechano et al., 2013) and alternative splicing events are widespread (Kahles et al., 2018), the set of post-transcriptional regulatory elements such as RNA-binding protein (RBP) binding sites, uORFs (Cheng et al., 2018; Tresenrider et al., 2021), or TOP motifs could be widely used in different biological contexts to link transcription and translation and direct the translational potential of an mRNA by the precise 5’ transcript sequence.

## Results

To systematically monitor the landscape of the full-length transcriptome, we performed nanopore long-read RNA sequencing of *in vivo* mouse epidermal stem cells and cultured squamous cell carcinoma cells (SCC^c^) to exemplify distinct biological contexts of the mouse skin. First, we sorted interfollicular epidermal stem cells (EpSCs) of P60 (postnatal day 60) adult mice by a rapid magnetic-activated cell sorting protocol using anti-stem cell antigen-1 microbeads (*Sca-1*, encoded by the *Ly6a* gene) (Hu et al., 2018; Joost et al., 2016). SCC^c^ were obtained from a previously established tumor allograft model, driven by oncogenic *HRAS*^*G12V*^ in combination with loss of the TGFβ receptor II, which rapidly form invasive squamous cell carcinomas when injected into *Nude* mice (Yang et al., 2015). As a reference, we used wild-type keratinocytes (Figure 1A).

**Figure 1.**
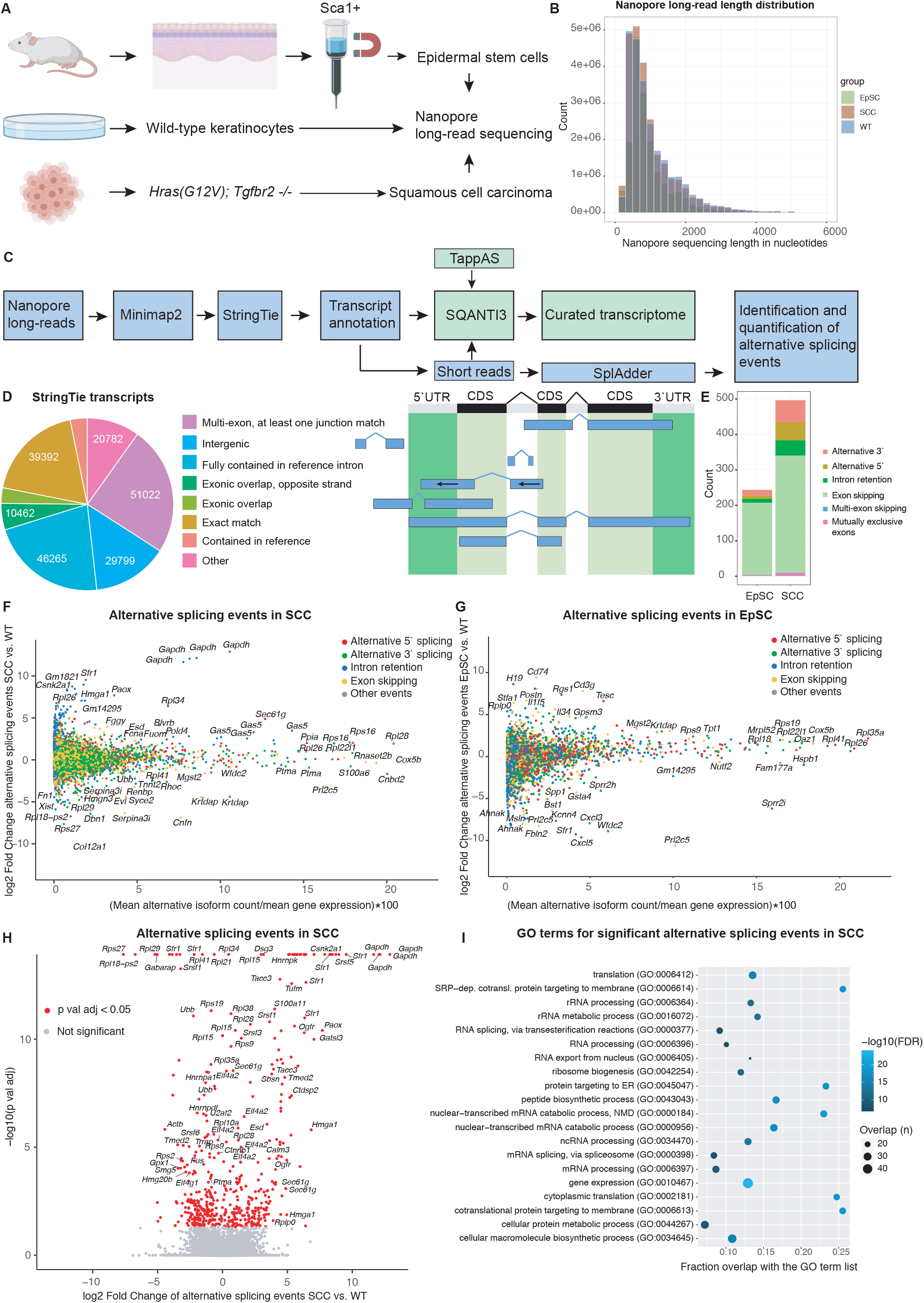
A. Experimental outline for the isolation of Sca-1+ epidermal stem cells, wild-type keratinocytes and cultured squamous cell carcinoma cells (SCC^c^) used for nanopore long-read sequencing. B. Read length distribution for the nanopore long-read sequencing data set of epidermal stem cells (EpSC), wild-type keratinocytes (WT), and squamous cell carcinomas (SCC^c^). The mean read length was between 943-1035 bp (EpSC: 943 bp, WT: 1035 bp, SCC^c^: 994 bp). C. Bioinformatic pipeline for the processing of the nanopore long-read sequencing data to identify and quantify alternative isoforms in epidermal stem cells and squamous cell carcinomas. D. Quantification and categorization of the total transcript numbers identified by StringTie using the nanopore long-read sequencing data of Sca1+ epidermal stem cells (EpSCs), wild-type keratinocytes, and squamous cell carcinomas (SCC^c^). Right panel shows examples for the different categories as defined by StringTie for transcript identifications. E. Numbers and categories of significantly changed isoforms in epidermal stem cells (EpSC) and squamous cell carcinomas (SCC^c^), compared to wild-type keratinocytes, as identified by the SplAdder pipeline. F-G. The landscape of alternative isoforms in squamous cell carcinomas or epidermal stem cells compared to wild-type keratinocytes using SplAdder, which quantifies and tests alternative splicing events. Color-coded are alternative 5’ splicing, 3’splicing, intron retention and exon skipping events. The x axis shows the alternatively spliced isoforms as a percentage of total gene expression. H. Volcano plot showing the significant events (red) and fold changes of splicing events in squamous cell carcinomas (SCC^c^) compared to wild-type keratinocytes (WT). I. Gene expression (GO:0010467) is the top gene ontology (GO) term for alternatively spliced genes in squamous cell carcinomas (SCC^c^) and is mostly driven by ribosomal proteins and splicing factors. GO term analysis shows the top 20 GO term hits (alphabetically ordered) with their false discovery rate (FDR, blue tone) and overlap with the GO term gene list in numbers (size of circle) and fraction (x axis).

In total, we generated 158.1 million reads with a mean read length between 943-1035 bp (Figure 1B). For transcriptome *de novo* assembly, we first used StringTie (Pertea et al., 2015), which allows accurate reconstructions and quantitation of genes and transcripts. Comparing the long-read sequences to the reference mouse genome, we identified a total of 210’630 transcripts, including 39’392 transcripts with an exact match of the intron chain to the reference genome (Figure 1C-D). For approximately half of the genes, we detected only one transcript, with another ∼35% of genes containing either two or three transcripts, while the remaining ∼15% had four or more transcripts (Figure supplement 1A). In addition, to classify and characterize the long-read transcript sequences, we used SQANTI3 to first map previously published matched short-read RNA-seq data (Sendoel et al., 2017) and to filter out transcripts that were not well-supported by the short-read data set, resulting in a curated annotation (Figure 1C).

To compute the occurrence of alternative isoforms and their differential expression in EpSCs and SCC^c^ versus wild-type keratinocytes, we exploited the SplAdder toolbox (Kahles et al., 2016) (Figure 1E-H). Using maximally stringent SplAdder confidence parameters, we found a total of 496 and 243 significantly changed isoform events in squamous cell carcinomas and epidermal stem cells, respectively. The identified alternative isoform events mainly fall into the categories of 5’ and 3’ alternative splice sites, exon skipping, and intron retention (Figures 1E). While in EpSCs, exon skipping events were clearly prevailing, SCC^c^ additionally exhibited 50 alternative 5’ and 63 alternative 3’ splicing events (Figure 1E). The alternative splicing events were observed across a wide range of relative isoform abundances compared to the corresponding gene expression in SCC^c^ and EpSCs (Figure 1F-G). Genes with significant alternative splicing events in SCC^c^ were enriched in ribosomal proteins and splicing factors (Figure 1H-I) and included a total of 23 ribosomal protein genes of the 40*S* and 60*S* ribosomal subunits (Figure 2A-B, Figure supplement 1B-C). Of note, many of these ribosomal proteins are located on the surface of the ribosome and include ribosomal proteins such as *Rpl38*, previously implicated in preferential translation of specific subpools of mRNA (Kondrashov et al., 2011). Together, our analyses document the landscape of alternative transcript isoforms and suggest that in squamous cell carcinomas, a small cohort of translation-related genes is differentially spliced.

**Figure 2.**
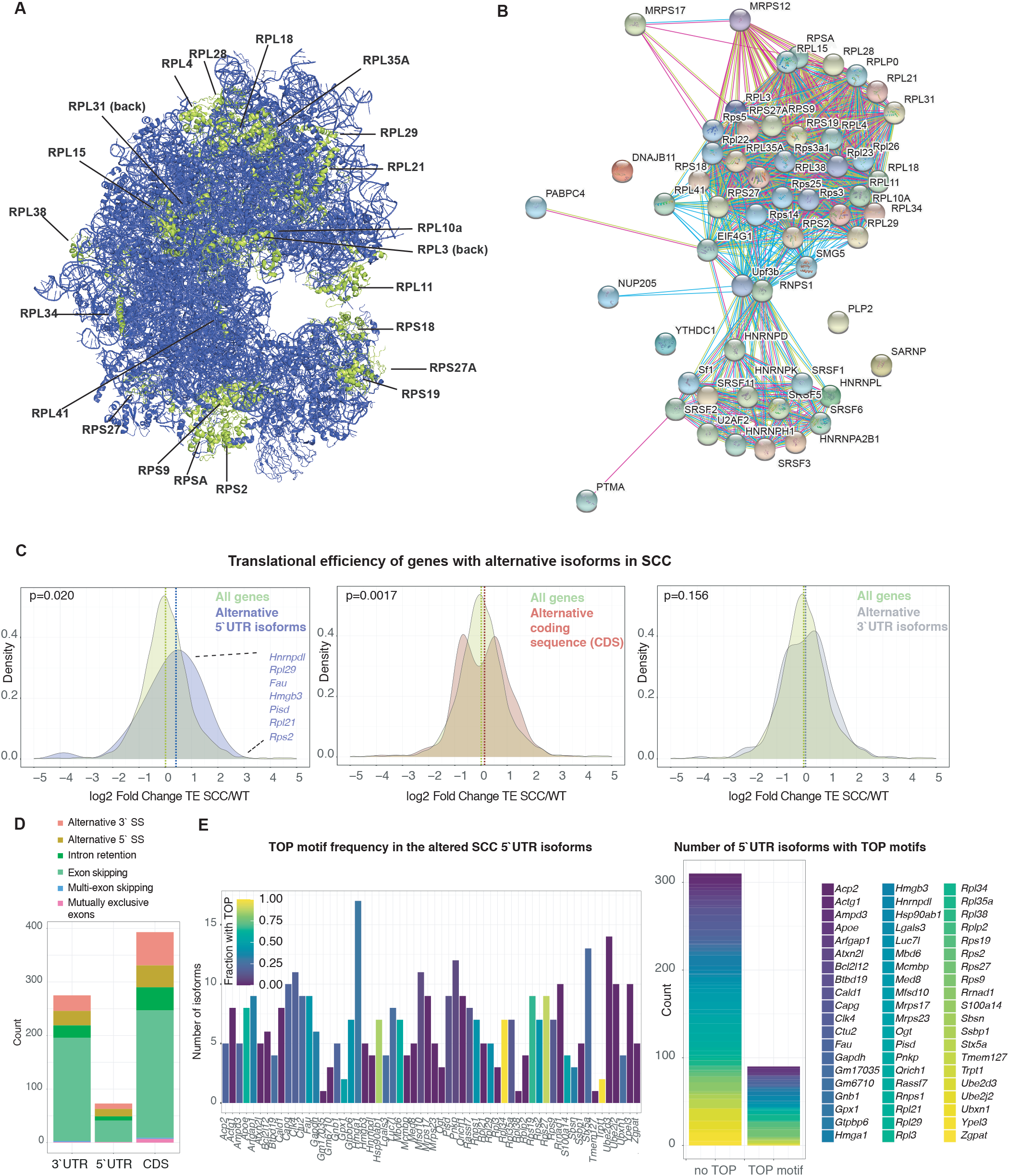
A. Localization of ribosomal proteins on the human 80*S* ribosome with significantly changed alternative splicing events of their encoding mRNAs in squamous cell carcinomas. Note that *Rplp0* and *Rplp2* were not included in the structure. B. STRING interaction network analysis for the alternatively spliced genes in the GO term gene expression (GO:0010467) in squamous cell carcinomas compared to wild-type keratinocytes. C. Translational efficiency (ribosome profiling reads divided by RNA-seq reads) of genes with differential alternative splicing events in the 5’UTR of squamous cell carcinomas (SCC^c^) is increased. Fold change in translation efficiency (TE) was computed for all genes or genes with significant alternative splicing events in the 5’UTR, coding sequence (CDS) or 3’UTR. P-values indicate a two-sample Kolmogorov-Smirnov test comparing the TE distribution of genes with alternative splicing events to all genes. D. Numbers and categories of significantly changed isoforms in squamous cell carcinomas (SCC^c^) that alter either the 5’UTR, the coding sequence (CDS) or the 3’UTR. E. Most genes with differential 5’UTR isoforms in squamous cell carcinomas (SCC^c^) express a set of transcripts that contain or exclude TOP motifs. StringTie transcripts and their 5’UTR sequences were assessed for the presence of TOP motifs as defined by a C (within the first 4 nucleotides) and an unbroken series of 4-16 pyrimidines. Left panel, colors indicate the fraction of StringTie transcripts that contain a TOP motif. Right panel, number of isoforms in the subset of genes with differential 5’UTR isoforms in SCC^c^ that contain a TOP or do not contain a TOP motif.

Given that the 5’UTR is a critical determinant of mRNA translation rates (Hinnebusch et al., 2016), we next focused on the alternative 5’UTR isoforms in SCC^s^ and asked how these 5’UTR isoforms affect translational efficiencies (TE). To determine which regions of the transcripts are affected by the alternative isoform usage, we further sub-grouped the significant alternative splicing events in SCC^c^ and found that 73 and 275 events altered the 5’UTRs and 3’UTR sequences, respectively. The identified alternative isoform events were mainly resulting from 5’ and 3’ alternative splice sites, intron retention, and exon skipping (Figure 2D). We then exploited a previously published ribosome profiling and short-read RNA-seq data set carried out in the identical wild-type keratinocyte and *Hras*^*G12V*^ ; *Tgfbr2 null* squamous cell carcinoma lines to compute differential translational efficiencies as a measure of protein synthesis rates per mRNA molecule (Sendoel et al., 2017). Of note, translational efficiency was assessed at the gene level. Therefore, low abundant isoforms with altered TEs would not be detected in the overall TE changes. As expected, when differential isoform usage between SCC^c^ and wild-type keratinocytes occurred in the 3’UTR, overall translational efficiencies of the altered genes were unaffected and exhibited a similar overall TE distribution compared to all genes (Figure 2C-D). In contrast, we found that the genes that exhibited significant alternative 5’UTR events resulted in a significant shift with an overall increase in translational efficiencies (Figure 2C, median log2 fold change 0.23 vs. 0.011 in 5’UTR isoform genes vs. all genes, p = 0.020, two-sample Kolmogorov-Smirnov test). Furthermore, the cohort of genes with alternative events in the CDS showed also a significant shift, however, with a bimodal distribution that did only slightly alter translational efficiencies (Figure 2C-D, median log2 fold change 0.13 vs. 0.011 in CDS isoform genes vs. all genes). These data suggest that squamous cell carcinomas differentially express alternative 5’UTR isoforms of a small cohort of genes to overall increase their protein synthesis rates.

mRNAs that encode translation factors typically possess 5’ terminal oligopyrimidine (TOP) motifs, which are essential for the coordinated translation of the family of TOP mRNAs (Philippe et al., 2020). Translation of TOP motif-containing mRNAs is orchestrated by the mammalian target of rapamycin complex 1 (mTORC1) nutrient-sensing signaling pathway (Hsieh et al., 2012; Thoreen et al., 2012). TOP motifs are defined as a +1 cytidine (C) directly adjacent to the 5′ cap structure, followed by an unbroken series of 4-16 pyrimidines. To determine the number of TOP motifs in the alternative 5’UTR isoforms of SCC^c^, we extracted the *de novo* 5’ StringTie transcriptome sequences of the genes with altered 5’UTR isoforms and analyzed them for the occurrence of TOP motifs. We found that most genes expressing altered 5’UTR isoforms in SCC^c^ contain both non-TOP and TOP motifs (Figure 2E), indicating the possibility of a cell to toggle between mTORC1-dependent and mTORC1-independent translation.

To experimentally test how individual 5’UTR isoforms impact the translational efficiency of an mRNA, we next focused on two ribosomal protein transcripts, *Rpl21* and *Rpl29*, which surfaced in our SplAdder analysis and showed distinct 5’UTR representations in the long-read nanopore sequencing data as also highlighted by the sashimi plots (Figure 3A-B). *Rpl21* and *Rpl29* express several isoforms differing in their 5’UTR sequences but contain identical coding and 3’UTR sequences. In addition, both, *Rpl21* and *Rpl29*, showed increased overall translational efficiencies in SCC^c^ compared to wild-type keratinocytes (log2 fold change of 1.38 and 1.25). We revised the 5’UTR annotation according to our *de novo* assembled transcriptome and cloned six 5’UTR isoforms of *Rpl21* and nine *Rpl29* 5’UTR isoforms into Firefly luciferase constructs to express them in wild-type keratinocytes and SCC^c^ (Figure 3A-B). To calculate translational efficiencies, we then measured *Rpl-*5’UTR Firefly and control-5’UTR *Renilla* luciferase activities and performed quantitative real-time PCR to assess mRNA levels. In addition, to address mTORC1 dependency of TOP motif-containing reporters experimentally, we treated the cells with the mTOR inhibitor Torin 1. We found that SCC^c^ generally showed higher TEs and that the different 5’UTRs resulted in more than an order of magnitude altered translational efficiencies of the Firefly luciferase (16.9x SCC/WT *Rpl29-8*, Figure 3D). The expression level of several *Rpl21* and *Rpl29* 5’UTR isoforms was significantly changed in SCC^c^ as quantified by the short-read RNA-seq data and the Ensembl annotation (Figure 3C). Furthermore, analyzing the long-read data, SCC^c^ showed a higher percentage of transcripts that contain 5’UTRs that were very efficiently translated, as exemplified by *Rpl21-2* or *Rpl29-1, Rpl29-5*, and *Rpl29-7* (Figure 3D-E, rpkm indicated below the figures). These TOP and TOP-like motif-containing isoforms showed mTORC1-dependent translation (Figure 3D-E). In contrast, wild-type keratinocytes exhibited a higher percentage of transcripts with low TEs, such as *Rpl21-4* and *Rpl21-5* (Figure 3C-E), which contain TOP motifs but were unchanged by Torin 1 treatment in wild-type keratinocytes. Together, these data exemplify that cancer cells can switch between mRNA 5’UTR isoforms to encode identical proteins produced with vastly altered efficiencies.

**Figure 3.**
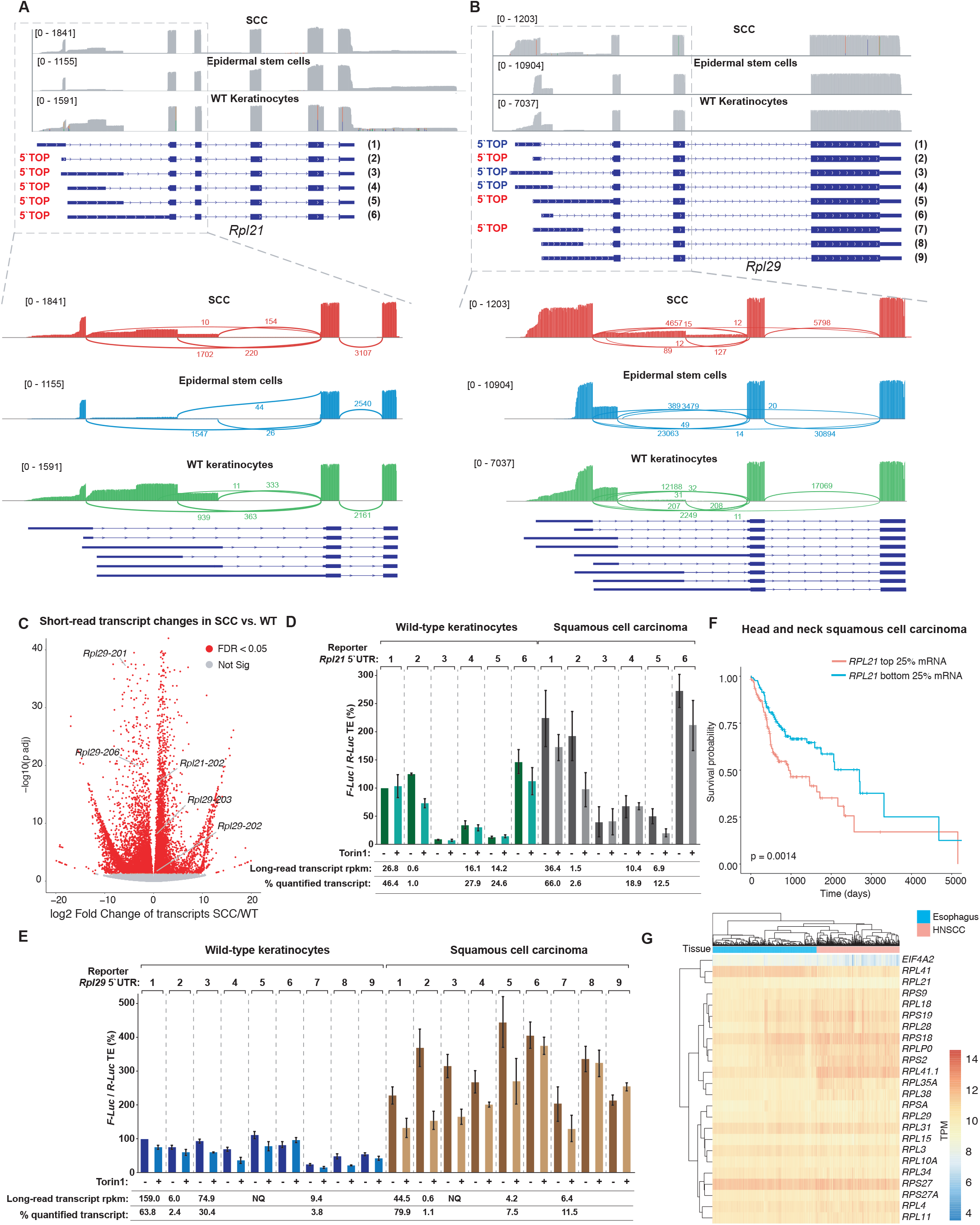
A, B. Representation of the nanopore long-read data set for *Rpl21* and *Rpl29* and their transcript annotation. Sashimi plots are shown below, highlighting the differential isoform usage of the transcript 5’UTRs. Red TOP indicates a transcript that contains a 5’ terminal oligopyrimidine motif, as defined by a C (within the first 4 nucleotides) and an unbroken series of 4-16 pyrimidines. Blue TOP indicates a transcript that contains a TOP-like motif as defined by an unbroken series of 4-16 pyrimidines, starting within the first 5 nucleotides. C. Volcano plot for DESeq2 differential transcript expression analysis in squamous cell carcinomas (SCC^c^) compared to wild-type keratinocytes (WT) reveals several *Rpl21/Rpl29* transcripts (Ensembl annotation) with significantly altered expression levels. The numbers next to the gene name refers to the Ensembl transcript ID. D-E. The different *Rpl21* and *Rpl29* 5’UTR isoforms show a wide range of translational efficiencies. Wild-type or squamous cell carcinomas (SCC^c^) were transfected with an *Rpl21* or *Rpl29* 5’UTR::Firefly-luciferase and a control 5’UTR::*Renilla*-luciferase plasmid and treated for 3 h with 500 nM of the mTORC1 inhibitor Torin 1 (+) or DMSO (-) before harvesting. The numbers refer to the transcript isoform numbers in Figure 3A. Below the graph are the long-read sequencing transcript rpkm (read per kilobase per million mapped reads) quantifications of the StringTie transcripts (if quantified by StringTie, otherwise empty field) and the percentage of the overall rpkm. NQ: StringTie transcript was detected and quantified in one condition but not the other. F. Increased *Rpl21* mRNA levels correlate with shorter overall survival of head and neck squamous cell carcinoma (HNSCC) patients. *Rpl21* top and bottom quartile mRNA expression in TCGA HNSCC patients’ samples (n=519). Cox regression hazard ratio 1.4629, p-value 0.00228. G. mRNA expression levels of the differentially spliced translation-related genes in TCGA head and neck squamous cell carcinoma (HNSCC) patients’ samples (n=519) and non-diseased esophageal tissue (non-diseased pharynx samples were not available).

## Discussion

Transcription and translation together determine ∼90% of cellular protein abundance (Schwanhäusser et al., 2011) and it is an intriguing question to what extent these two processes can be directly coupled in different biological contexts. In this short report, we carry out nanopore long-read sequencing to document the UTR isoform landscape of *in vivo* epidermal stem cells and cultured squamous cell carcinoma cells. We show that a small cohort of 5’UTR isoforms, differentially expressed in SCC^c^, can alter the translational efficiency of the corresponding coding sequence. Our analyses revealed that SCC^c^ are able to switch across these 5’UTR isoforms of mostly translation-related genes to expose or mask 5’terminal oligopyrimidine (TOP) motifs, which in the case of *Rpl21* and *Rpl29* resulted in markedly changed protein synthesis rates. Notably, human *RPL21* but not *RPL29* levels correlated with overall survival in head and neck squamous cell carcinomas, suggesting that such 5’UTR isoform switches may also be relevant for disease progression in cancer patients (Figure 3F-G). Nevertheless, given the relatively small cohort of 5’UTR isoform switches, our observations also suggest that the network of translational regulators is genome-wide still the main factor determining protein synthesis rates.

5’UTR isoform switches have emerged as an elegant non-canonical mode of gene expression regulation (Chen et al., 2017; Cheng et al., 2018; Chia et al., 2017; Tresenrider et al., 2021). Our findings add yet another simple mechanism to the 5’UTR landscape and suggest that exposing or masking TOP motifs from the 5’UTR can markedly alter the translational efficiency of the mRNA coding sequence. Translational efficiency changes could be achieved by relatively small changes in the exact transcription start site (TSS) to modify the presence of 5’ TOP motifs to set the dependency on mTORC1-dependent nutrient sensing to specifically enhance the translation of the family of TOP mRNAs (Hsieh et al., 2012; Thoreen et al., 2012). Given that alternative splicing and TSSs are widespread (Kahles et al., 2018; Pelechano et al., 2013), switching between 5’UTR isoforms with different sets of post-transcriptional regulatory elements such as uORFs, RBP binding motifs, or TOP motifs could be used in various biological contexts to directly encode the mRNA’s translational potential in its 5’UTR.

## Figure legends

**Figure supplement 1**.

A. Number of transcripts and coding sequences per gene as identified by the tappAS pipeline.

B. Gene expression is the top gene ontology (GO) term for alternatively spliced genes also in epidermal stem cells. GO term analysis shows the top 20 GO term hits (alphabetically ordered) with their false discovery rate (FDR, blue tone) and overlap with the GO term gene list in numbers (size of circle) and percentages (x axis).

C. Enrichment score in Gene Set Enrichment Analysis (GSEA) analysis for the GO term translation for the genes differentially spliced in SCC^c^. GO term analysis shows the top 20 GO term hits (alphabetically ordered) with their false discovery rate (FDR, blue tone) and overlap with the GO term gene list in numbers (size of circle) and fraction (x axis).

## Material & Methods

### Nanopore long-read RNA sequencing

RNA was isolated using TRIzol LS (Thermo Fisher, 10296010) and the Direct-zol RNA Miniprep Kit (Zymo Research, R2050). The samples were then used for library preparation following the manufacturer’s protocol of SQK-PCS109 (Oxford NANOPORE Technologies, Oxford Science Park, UK). Briefly, total RNA was polyA enriched using oligo dT, annealed with primers, and reverse transcribed. Following template switching, PCR with rapid attachment primers was performed and rapid 1D sequencing adapters were attached. The library was then sequenced on PromethION (Oxford NANOPORE Technologies, UK).

### Isolation of adult mid-telogen EpSCs

Female C57BL6 mice were obtained from Janvier. EpSCs were isolated from telogen back skin collected from P56-60 mice, as previously described (Nowak and Fuchs, 2009) with the following changes: Fat and muscle tissue was removed from back skin using a scalpel. The skin was incubated in 0.5% Trypsin-EDTA (10X; Gibco,15400054) for 25 minutes at 37°C on an orbital shaker. A single-cell suspension was then obtained by scraping the skin with a scalpel followed by neutralizing the trypsin by adding 1X PBS buffer containing 2% chelexed FBS (PBS-FBS(-)) (Gibco; 10010-015). The resulting cell suspension was then filtered through 70-µm and 40-μm cell strainers (Corning; 431750, 431751) and spun down. EpSCs were isolated using magnetic-associated cell sorting using Anti-Sca-1 microbeads (Miltenyi Biotec, 130-106-641) and a MACS MultiStand system (Miltenyi Biotec) together with MS columns (Miltenyi Biotec, 130-042-201). The resulting SCA-1^+^ EpSCs were spun down and resuspended in Trizol LS (Thermo Fisher, 102960-10). RNA concentration was determined using the Qubit™ RNA BR assay kit (Invitrogen, Q10210). All animal procedures were approved by the Veterinary Office of the Canton of Zurich, Switzerland (License ZH233/2019).

### *In vitro* cell culture experiments

Newborn, primary mouse epidermal keratinocytes from wild-type mice were cultured on 3T3-S2 feeder layer in 0.05 mM Ca2+ E-media supplemented with 15% serum (Nowak and Fuchs, 2009). *HrasG12V; Tgfbr2* knockout cell line was previously generated (Yang et al., 2015). Cell lines were cultured in E medium with 15% FBS and 50 mM CaCl2 (Nowak and Fuchs, 2009). Cell lines were tested for mycoplasma infection every 3 months.

### Luciferase assays

0.25×10^6^ cells were plated in a total volume of 2 ml 0.05 mM Ca2+ E-media in 6-well plates 24 h before transfection. The transfection mixtures contained 1.5 µg Rpl29/Rpl21-F-Luc and 0.5 µg control R-Luc plasmid DNA. All transfections were performed using Lipofectamine 2000. Two hours after transfection cells were treated with 500 nM Torin 1 or DMSO and harvested five hours after transfection. Cells were briefly washed with PBS and lysed in passive lysis buffer. *Firefly* and *Renilla* luciferase activities were measured at room temperature using the Dual-Luciferase reporter assay system (Promega, E1980) and a Tecan Infinite M1000Pro instrument.

### Quantitative real-time PCR

For RNA extraction, cells were resuspended in TRIzol LS (Thermo Fisher, 10296010) and extracted using chloroform. The aqueous phase was then precipitated in isopropanol, pelleted, and resuspended in H2O. For cDNA synthesis 0.5 µg total RNA was mixed with 0.5 µg of random hexamer primers (N6) and denatured at 72 °C for 5 min. Subsequently, a reaction mixture containing 1x SSIII RT buffer, 1 mM dNTPs, 5 mM DTT, and 0.5 µl SSIII (Invitrogen, 18080093) was added to reach a total volume of 20 µl. The RT reaction was performed at 55°C for one hour and inactivated for 10 min at 70 °C. The qPCR was performed in a final concentration of 1x iTaq Universal SYBR Green Supermix (Biorad, 1725121), 0.4 µM primer each, and 1 µl of the cDNA in a total volume of 10 µl.

### Bioinformatic analyses

Sample processing and analysis were performed on an Ubuntu 18.04.5 cluster with 32 cores and 128GB RAM. Standard parameters were used for all software unless stated otherwise, Python version is 3.8.6 unless indicated otherwise.

### Processing of nanopore long-read data

The long-read sequencing data were processed by Nextflow nanoseq v1.1 (Patel et al., 2020) and a custom pipeline. The quality of the raw fastq files and the sequencing was assessed by the FastQC and NanoPlot (De Coster et al., 2018) programs. Alignment to the mouse reference genome Gencode GRCm38 version M25 was carried out by minimap2 which can do both spliced and unspliced alignment. Custom transcriptome reference assembly and expression quantification were performed by StringTie2 (Pertea et al., 2015). Furthermore, the transcriptomes were merged and compared by GFFcompare v.0.11.2 (Pertea and Pertea, 2020). Nanopore long-read data are available on the this paper’s GEO.

### Processing of short-read RNA-seq data

We utilized the Nextflow rnaseq pipeline (Patel et al., 2020) and custom scripts to process the short-read RNA-seq data. The quality of the fastq files was assessed by the FastQC program. TrimGalore was used to perform quality and adapter trimming of the sequencing data. The reads were aligned to the GRCm38 reference by STAR v2.6.1 (Dobin et al., 2013). Transcriptome quantification was carried out by Salmon (Patro et al., 2017). BigWig files were created by BEDTools for visualization of coverage tracks. Furthermore, MultiQC was used to carry out quality control of all the analysis pipelines.

### Differential Expression

The raw gene and transcript level counts obtained were processed by DESeq2 to call the differentially expressed (DE) genes between various conditions (Love et al., 2014). Hierarchical clustering and PCA plots after variance stabilizing transformation (vst) normalization of the top 500 most variable genes were used to detect any outlier samples. First, the count data were normalized by the median of ratios method. Next, the dispersion or biological variance was estimated. A generalized linear model was fitted for each gene to detect differentially expressed (DE) genes. The p-values obtained by the Wald test were corrected by the Benjamini–Hochberg multiple testing procedure. DE genes with FDR cut-off <0.05 were used for further analysis.

### Processing of ribosome profiling samples

Ribosome profiling samples were processed as described in the short-read RNA-seq section above, with the addition of a filtering step after the adapter trimming. Only reads not aligning to a merged transcriptome consisting of ribosomal, mitochondrial, or tRNA (Chan and Lowe, 2016) sequences using Bowtie2 v2.4.1 (Langmead et al., 2019) were processed further.

### Ribosome profiling and short-read RNA sequencing samples

Ribosome profiling and short-read RNA sequencing data were used from a previously published study (Sendoel et al., 2017) with matched wild-type keratinocyte and *Hras*^*(G12V);*^ *Tgfbr2* null squamous cell carcinoma samples, which are available on the GEO GSE83332. We used the following samples:

Wild-type keratinocytes: S11, S13 and S14 (Ribosome profiling, GSM2199591, GSM2199593, GSM2199594) and S27 and S28 (short-read RNA-seq, GSM2199607, GSM2199608).

Squamous cell carcinomas: samples S23, sample 197 and S24 (Ribosome profiling, GSM2199603, this paper’s GEO, GSM2199604) and samples 211 and 212 (short-read RNA-seq, this paper’s GEO).

Epidermal stem cells: samples S7 and S8 (short-read RNA-seq, GSM2199587, GSM2199588), which are P4 epidermis samples enriched for epidermal stem cells. Note that ribosome profiling data were not available for P60 Sca-1+ epidermal stem cells.

### Curated transcriptome

For the curated transcriptome, long read isoform quantification and characterization were also performed by SQANTI3 (Tardaguila et al., 2018), using short-read RNA-seq samples in a sequential manner. First, the whole StringTie assembled transcriptome was used to quantify isoforms and subsequently subjected to the SQANTI3 filtering to remove not well-supported junctions and antisense transcripts passively produced by the ONT1D procedure. The obtained filtered annotation was used to create a new filtered transcriptome and re-map samples to increase mapping rates and counts. Functional annotations were transferred from a pre-computed tappAS annotation file (De La Fuente et al., 2020) based on PacBio sequencing by facilitating the IsoAnnotLite (v2.6) option of SQANTI3.

### Translational efficiency

Genomic reads of RNA-seq and ribosome profiling samples were quantified by Plastid v0.4.7 (Dunn and Weissman, 2016) (Python 2.7) over extracted exon and coding sequences (CDS) regions, respectively, as outlined previously (Sendoel et al., 2017). Translational efficiency (TE) was computed in R v4.0.2 using the LRT-test of the DESeq2 package and full/reduced models as suggested by the plastid authors (Dunn and Weissman, 2016) for genes that showed rpkm (read per kilobase per million mapped reads) > 1 in RNA-seq samples. For comparison with the SplAdder gene lists (Figure 2C), we included TE calculations with a base mean > 25. Note that raw counts were used for DESeq2 and not rpkm, therefore differences in rpkm between samples are normalized out.

### Differential isoform events

Alternative splicing events were computed bySplAdder v2.4.3 (Kahles et al., 2016) using maximum level 3 confidence parameters and the StringTie annotations and genome mapped short-read RNA-seq samples. Results were subjected to a threshold filtering, requiring an adjusted p-value < 0.05.

### Pathway Enrichment

Selected genes were analyzed for gene ontology (GO) pathway enrichment using GSEA pre-ranked method which enables the analysis of up- and down-regulated genes simultaneously. This approach significantly improves the sensitivity of the gene set enrichment analysis. The overrepresentation analysis for the pathway enrichment was carried out by EnrichR and custom scripts.

### TCGA and GTEx data analysis

TCGA and GTEx data were obtained for HNSCC and esophagus from UCSC Xena Toil (xena.ucsc.edu). Survival analysis for TCGA HNSCC was performed by the survival and survminer library in R. A Cox Proportional Hazards regression model was used to fit the gene expression to survival to obtain the Hazard Ratio. Kaplan-Meier analysis was performed on groups with lower and upper quartiles of gene expression and the p-values were computed by a log-rank test.

## Acknowledgments

This project was supported by an SNSF Professorship grant (grant number 176825).

---

**Extended Data Figure 1.**
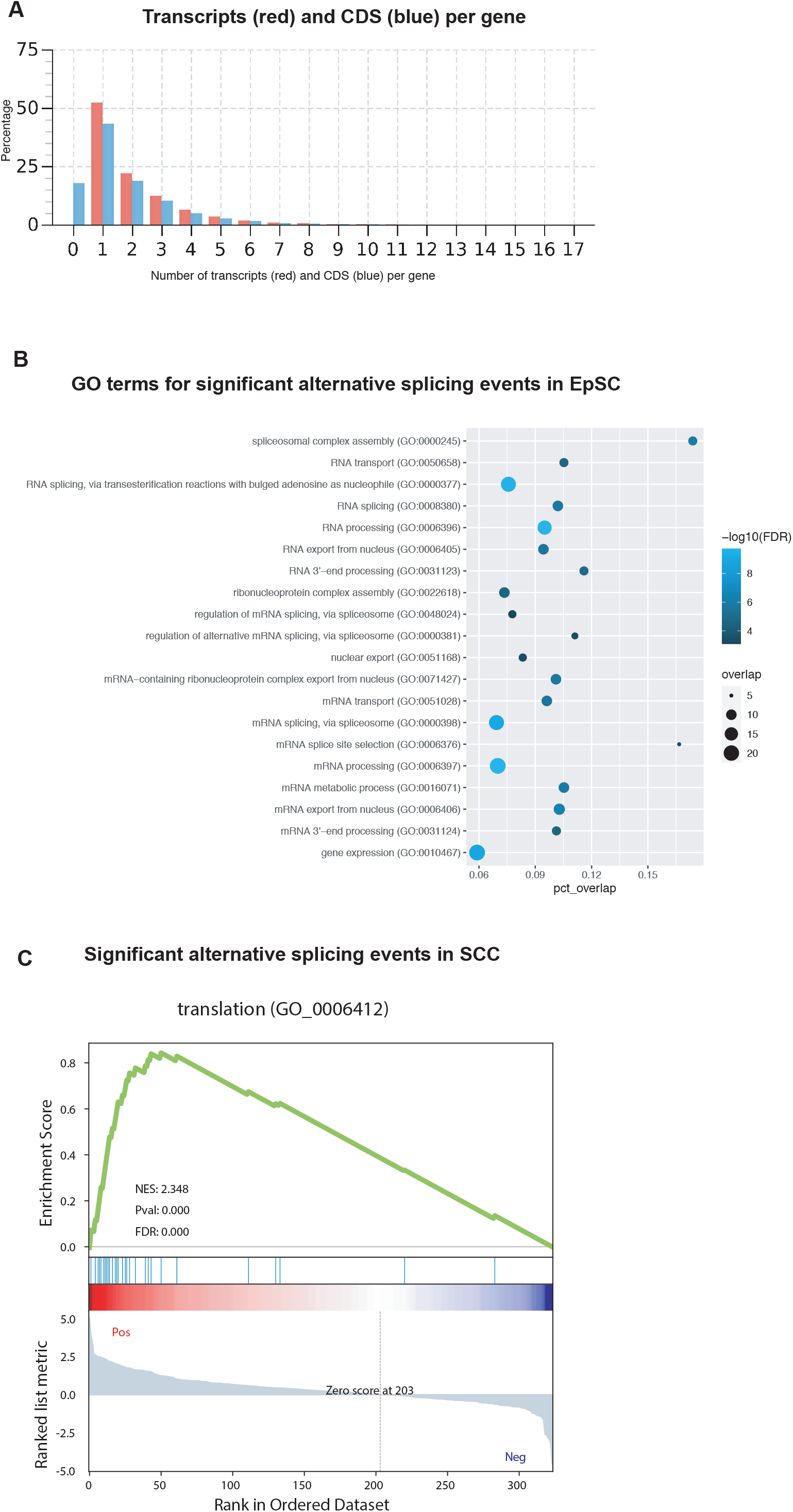

